# EP3 signaling is decoupled from regulation of glucose-stimulated insulin secretion in β-cells compensating for obesity and insulin resistance

**DOI:** 10.1101/2020.07.10.197863

**Authors:** Michael D. Schaid, Jeffrey M. Harrington, Grant M. Kelly, Sophia M. Sdao, Matthew J. Merrins, Michelle E. Kimple

**Author notes:** Co-corresponding authors: Michelle E. Kimple, Department of Medicine, Division of Endocrinology, Diabetes, and Metabolism, Medical Foundation Centennial Building, 1685 Highland Avenue, University of Wisconsin-Madison 53705, Matthew J. Merrins, Department of Medicine, Division of Endocrinology, Diabetes, and Metabolism, Medical Foundation Centennial Building, 1685 Highland Avenue, University of Wisconsin-Madison 53705.

## Abstract

Of the β-cell signaling pathways altered by non-diabetic obesity and insulin resistance, some are adaptive while others actively contribute to β-cell failure and demise. Cytoplasmic calcium (Ca^2+^) and cyclic AMP (cAMP), which control the timing and amplitude of insulin secretion, are two important signaling intermediates that can be controlled by stimulatory and inhibitory G protein-coupled receptors. Previous work has shown the importance of the cAMP-inhibitory EP3 receptor in the beta-cell dysfunction of type 2 diabetes. To examine alterations in β-cell cAMP during diabetes progression we utilized a β-cell specific cAMP biosensor in tandem with islet Ca^2+^ recordings and insulin secretion assays. Three groups of C57BL/6J mice were used as a model of the progression from metabolic health to type 2 diabetes: wildtype, normoglycemic *Leptin*^*Ob*^, and hyperglycemic *Leptin*^*Ob*^. Here, we report robust increases in β-cell cAMP and insulin secretion responses in normoglycemic *Leptin*^*ob*^ mice as compared to wild-type: an effect that was lost in islets from hyperglycemic *Leptin*^*ob*^ mice, despite elevated Ca^2+^ duty cycle. Yet, the correlation of EP3 expression and activity to reduce cAMP levels and Ca^2+^ duty cycle with reduced insulin secretion only held true in hyperglycemic *Leptin*^*Ob*^ mice. Our results suggest alterations in beta-cell EP3 signaling may be both adaptive and maladaptive and define β-cell EP3 signaling as much more nuanced than previously understood.

## Introduction

Type 2 diabetes (T2D) is associated with obesity and insulin resistance, yet not all insulin resistant individuals are diabetic. Pancreatic β-cells can compensate via increased glucose sensitivity, mass, or both, resulting in circulating insulin levels sufficient to maintain normoglycemia. The second messenger cyclic AMP (cAMP) plays a crucial role in β-cell compensation, and while many T2D therapies aim to increase β-cell cAMP levels, the efficacy of these therapies is not sufficient for many patients (1). Previous work by our group and others identified enhanced signaling through the prostaglandin EP3 receptor (EP3), a cAMP-inhibitory G protein-coupled receptor (GPCR) encoded by the *PTGER3* gene, in islets isolated from T2D mice and humans as compared to non-diabetic controls (2-6). The most abundant natural ligand of EP3 is prostaglandin E_2_ (PGE_2_), an eicosanoid derived from arachidonic acid whose islet production is similarly increased in the pathophysiological context of T2D (2,6-10). EP3 agonists reduce, while EP3 antagonists potentiate glucose-stimulated insulin secretion (GSIS) of islets from T2D mice and humans, while having little to no effect on islets from wild-type or non-diabetic subjects (2,8,10).

In β-cell lines and pancreatic islets, EP3 is specifically coupled to the G_i/o_ subfamily of inhibitory G proteins to suppress cAMP production and GSIS (2,11-16). Yet, the role of EP3 and its natural ligands in T2D remains controversial. In the context of obesity, PGE2 and other arachidonic acid metabolites have tissue-specific beneficial effects on inflammation and insulin resistance, and EP3 knockout mice are more prone to metabolic dysfunction after high-fat-diet feeding (17-19). Furthermore, evidence exists for agonist- and cAMP-independent effects of EP3 activity on cellular function and secretion processes (20-22). In this work, we examined the relationship of EP3 expression and signaling on β-cell cAMP homeostasis, Ca^2+^ influx, and GSIS during the progression from metabolic health, to β-cell compensation, and, finally, T2D. Using islets isolated from C57BL/6J mice, either wild-type or homozygous for the *Leptin*^*Ob*^ mutation, in combination with high-sensitivity, temporal imaging of β-cell cAMP levels and Ca^2+^ oscillations before and after receptor ligand stimulation, we find that increased EP3 expression correlates directly with the effects of a selective EP3 agonist on cAMP production and Ca^2+^ influx, but that EP3 signaling in compensating β-cells is uncoupled from regulation of GSIS.

## Results

### EP3 expression, but not PGE_2_ production, is associated with progression towards diabetes in *Leptin*^*Ob*^ mice

Using a random-fed blood glucose level of 300 mg/dl, two distinct populations of 10-12 week-old male C57BL/6J *Leptin*^*Ob*^ mice were identified, both as compared to wild-type C57BL/6J males – normoglycemic (NGOB) and hyperglycemic (HGOB) (Fig. 1A). These results are consistent with a previous report plotting the relationship between plasma insulin and plasma glucose levels in a population of C57BL/6J *Leptin*^*Ob*^ males (23). The mean random-fed blood glucose levels of wildtype (WT) and NGOB were nearly identical (214 mg/dl and 212 mg/dl, respectively), whereas that of HGOB mice was just over twice as high (438.9 mg/dl, p<0.0001). The mean islet insulin content was dramatically enhanced in NGOB as compared to WT islets, and trended lower in HGOB islets, consistent with T2D status (Fig. 1B). EP3 (*Ptger3*) mRNA abundance was significantly higher in islets from both NGOB and HGOB mice as compared to WT (Fig. 1C).

**Figure 1.**
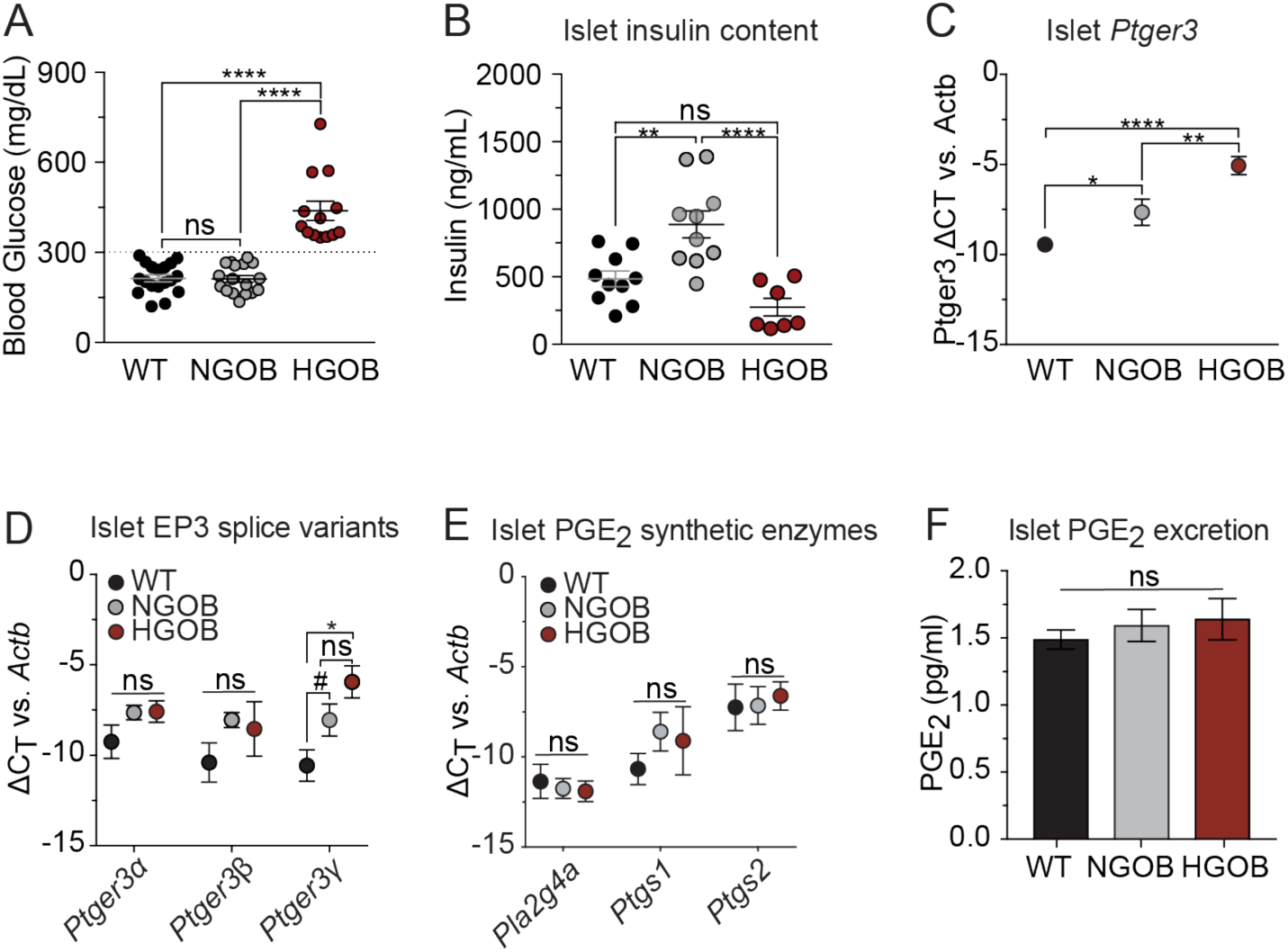
Expression of EP3 is associated with impaired insulin secretion independent of PGE_2_ production. (A) Blood glucose levels in C57BL/6J wild-type (WT), normoglycemic *Leptin*^ob^ (NGOB) and hyperglycemic *Leptin*^ob^ (HGOB) mice. N=13-18. (B) Islet insulin content. N=7-10. (C) Relative *Ptger3* transcript levels normalized to *Actb*. N=9-15. (D) Relative *Ptger3* isoform transcript levels normalized to *Actb*. N=4-7. (E) Selected PGE_2_ synthetic enzymes (*Pt2g4a, Ptgs1*, and *Ptgs2*) as normalized to *Actb*. N=4-7. (F) 24 h PGE_2_ excretion from cultured islets. N=6-8. In all cases, N=individual mice or mouse islet preparations, and data represent the mean ± SEM. Data was analyzed by one-way ANOVA with Tukey test post-hoc (A-E) or multiple t-test corrected with Holm-Sidak test post-hoc (G). ^#^, P=0.06; *P<0.05; **P<0.01; ***P<0.001; and ****P<0.0001. ns = not significant.

The mouse *Ptger3* gene encodes three EP3 splice variants, EP3α, EP3β, and EP3γ, that differ only in their cytoplasmic C-terminal tail, a region important for properties including G protein coupling and constitutive activity (12,24). Using isoform-specific qPCR primers, the mean EP3α and EP3β mRNA abundance was modestly elevated in both NGOB and HGOB islets; an effect that was not statistically-significant (Fig. 1D, *left* and *middle*). However, EP3γ mRNA abundance exhibited the same step-wise increases from WT to NGOB to HGOB as observed with total *Ptger3* expression (Fig. 1D, right). Neither NGOB nor HGOB mouse islets had any significant alteration in the mRNA abundance of the PGE_2_ synthetic enzymes important in the β-cell (2,10) as compared to WT controls (Fig. 1E). *Ptgs1* was the only mRNA with a statistically-significant increased abundance in NGOB or HGOB islets as compared to WT, while that of *Ptgs2*, the dominant isoform in β-cells (25), was unchanged. Consistent with a lack of upregulation of important PGE_2_ synthetic genes, islet PGE_2_ excretion was nearly identical among the three groups (Fig. 1F).

### Validation of the cAMP FRET assay and islet phenotypes

In this work, we utilized a high-affinity Exchange protein activated by cAMP (Epac)-based cAMP FRET sensor (Epac-S^H187^) (26), adenovirally-expressed under the control of the rat insulin promoter in order to confer β-cell specificity (27). A representative two-photon microscopy image of a transduced islet shows penetrance of the expression into the interior of the islet (Fig. 2A). Transduced islets perfused with imaging buffer showed a dramatic shift in fluorophore intensity in response to 3-isobutyl-1-methylxanthine (IBMX, 100 µM), a phosphodiesterase inhibitor (Fig. 2B, *left*), increasing the normalized and average FRET ratio (R470/530) (Fig. 2B, *middle and right)*.

**Figure 2.**
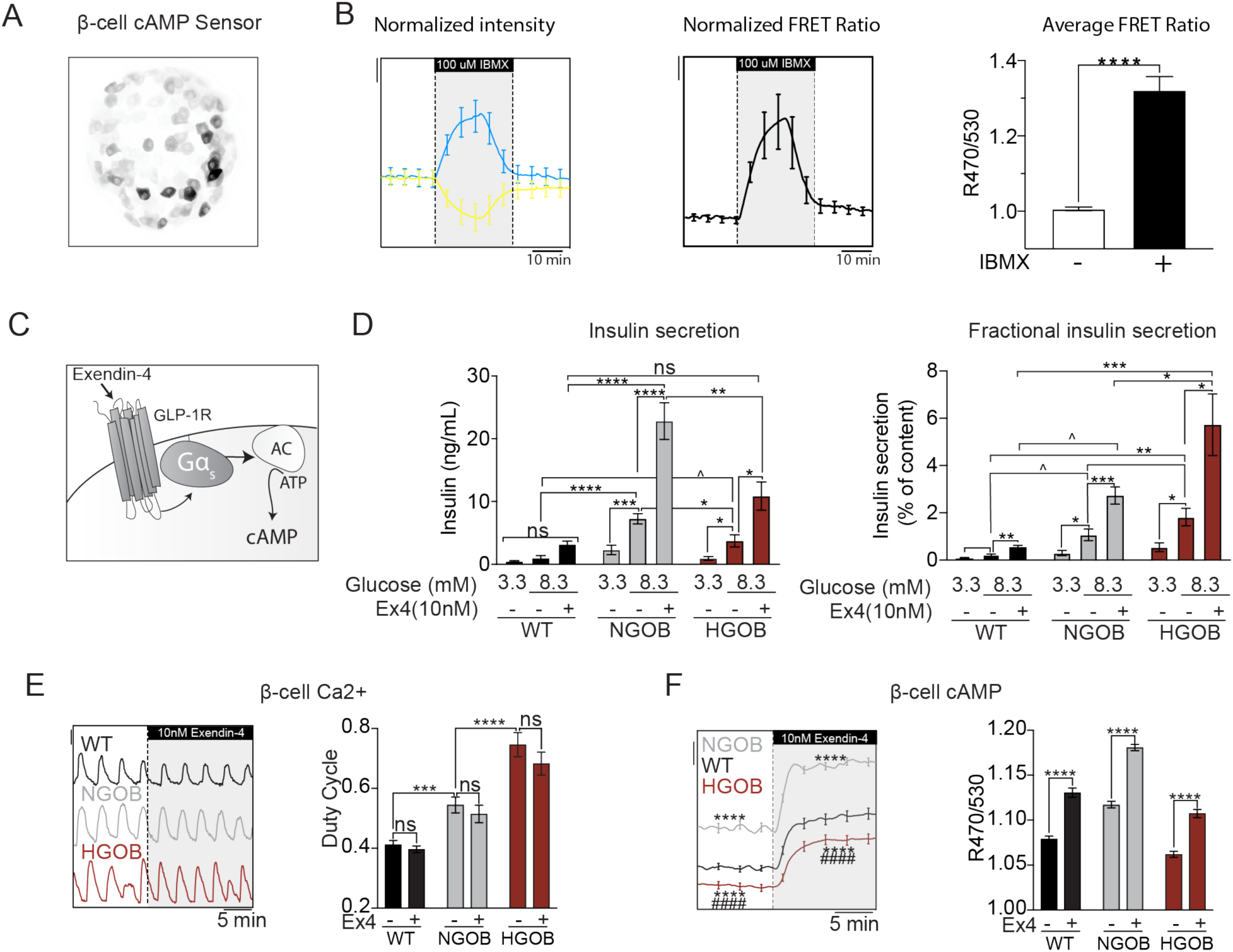
Validation of β-cell-specific cAMP FRET sensor and correlation of β-cell cAMP levels with islet pathophysiology. (A) 3D projection of WT islets expressing the β-cell specific cAMP biosensor (Epac-S^H187^) as recorded by two-photon microscopy. (B) Normalized fluorophore intensity (left), FRET ratio (middle) and average FRET ratio (right) from WT islets expressing cAMP biosensor treated with 100uM IBMX. N=7 islets. (C) Diagram of the GLP-1 receptor agonist Exendin-4 (Ex4) and its effects on cAMP production. (D) Glucose stimulated insulin secretion from WT, NGOB and HGOB islets treated with 3.3 mM glucose or 8.3 mM glucose ± 10 nM Ex4. Data is represented as both total insulin secreted (left) and fractional insulin secretion as a percent of total insulin content. N=7-10 mice. (E) Representative Ca^2+^ recordings and duty cycle quantification of WT, NGOB and HGOB islets treated with 9 mM glucose followed by 9 mM glucose +10 nM Ex4. (F) Representative average cAMP recordings and quantification of WT, NGOB and HGOB islets treated with 9 mM glucose followed by 9 mM glucose +10 nM Ex4. N=55-72 islets from 3 mice (NGOB and HGOB) and 90 islets from 6 mice (WT). Scale bar: FuraRed (Ca^2+^)= 0.01 (D) or, 0.2 (B) and 0.025 (E) R470/530 (cAMP). Error bars represent the standard error of the mean (SEM). Data was analyzed by one-way ANOVA or multiple student t-test with Tukey test post-hoc.*p<0.05, **p<0.01, ***p<0.001, ****p<0.0001. For panel E, ****p<0.0001 as compared to WT, #### p<0.0001 as compared to NGOB.

Exendin-4 (Ex4) is a stable GLP-1 receptor agonist that is known to potentiate GSIS by cAMP-mediated mechanisms. Therefore, Ex4 was used to both quantify the secretion phenotype of islets from WT, NGOB, and HGOB mice and confirm its relationship with β-cell cAMP levels. Consistent with their β-cell compensation phenotype, islets from NGOB mice secrete significantly more insulin in 8.3 mM glucose than 3.3 mM glucose, and their mean GSIS response is significantly higher than WT, whether represented as total insulin secreted or insulin secreted as a percent of content (Fig. 2D). Islets from HGOB mice secrete significantly less insulin in 8.3 mM glucose than NGOB, and their fractional GSIS response was higher (Fig. 2D), consistent with their T2D phenotype. All three groups responded strongly to exendin-4 (Ex4) to potentiate GSIS by approximately 2-fold over 8.3 mM glucose alone, whether represented as total or fraction of content (Fig. 2D). Imaging experiments performed in parallel with GSIS assays reveal that Ca^2+^ duty cycle (fractional time in the active phase of an oscillation)(28) was sequentially increased in NGOB and HGOB islets as compared to WT, but is unaffected by Ex4 in any group (Fig. 2E). In contrast, β-cell cAMP is strongly and significantly increased in NGOB islets as compared to WT and reduced in HGOB islets as compared to the other groups (Fig. 2F). In all three cases, consistent with GSIS results, Ex4 significantly increased cAMP levels (Fig. 2F).

### Endogenous agonist-dependent EP3 signaling does not contribute to β-cell compensation or dysfunction in B6J *Leptin*^*Ob*^ islets

Both PGE_2_ production and EP3 expression have previously been shown to contribute to β-cell dysfunction in the BTBR *Leptin*^*Ob*^ mouse model, a unique and severe model of T2D (2). In the current work, islet PGE2 synthetic gene expression and PGE_2_ excretion are both unchanged in NGOB or HGOB islets vs. WT (Fig. 1E,F), suggesting agonist-sensitive EP3 signaling does not contribute to either β-cell compensation or β-cell failure in this model. To confirm this at the molecular level, we utilized the EP3-specific antagonist, DG041 (10 nM), which blocks binding of endogenous PGE_2_ to EP3 (Fig. 3A). As expected, DG041 had no effect on GSIS (Fig. 3B), cAMP levels (Fig. 3C), or Ca^2+^ duty cycle (Fig. 3D).

**Figure 3.**
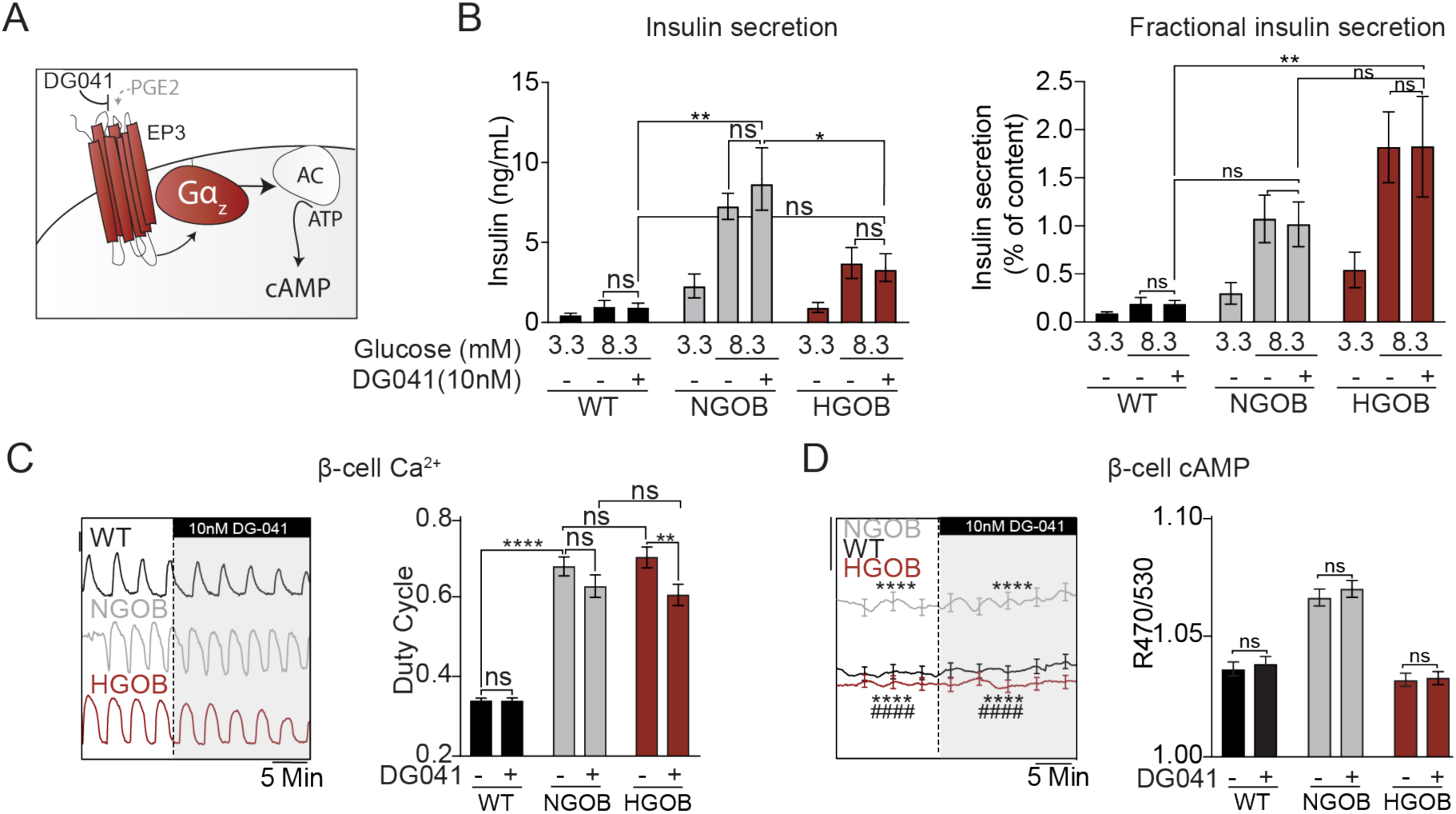
Agonist-sensitive EP3 signaling does not influence WT, NGOB, or HGOB β-cell cAMP production or function. (A) Diagram of the EP3 receptor antagonist DG041 and its effects on cAMP production. (B) Glucose stimulated insulin secretion from WT, NGOB and HGOB islets treated with 3.3 mM glucose or 8.3 mM glucose ± 10 nM DG041. Data is represented as both total insulin secreted (left) and fractional insulin secretion as a percent of total insulin content. N=7-10 mice. (C) Representative Ca^2+^ recordings and duty cycle quantification of WT, NGOB and HGOB islets treated with 9 mM glucose followed by 9 mM glucose +10 nM DG041. (D) Representative average cAMP recordings and quantification of WT, NGOB and HGOB islets treated with 9mM glucose followed by 9 mM glucose +10 nM DG041. In ‘C’ and ‘D’ N=55-72 islets from 3 mice (NGOB and HGOB) and 108 islets from 6 mice (WT). Scale bar: FuraRed (Ca2+)= 0.01 (B) or 0.025 (C) R470/530 (cAMP). Error bars represent the standard error of the mean (SEM). Data was analyzed by one-way ANOVA or multiple student t-test with Tukey test post-hoc.*p<0.05, **p<0.01, ***p<0.001, ****p<0.0001. For panel D, ****p<0.0001 as compared to WT, #### p<0.0001 as compared to NGOB.

EP3γ is the only splice variant that is significantly enhanced in NGOB or HGOB islets as compared to WT, and its expression follows the same pattern as total EP3 expression (Fig. 1C,D). Importantly, EP3γ is approximately 80% constitutively active, meaning that it does not require PGE_2_ binding to signal downstream (29,30). Yet, as it remains partially agonist-sensitive, we can utilize the EP3-selective agonist, sulprostone, to explore its downstream effects (Fig. 4A). Consistent with previous reports using islets from lean and/or non-diabetic mice and humans (2,10,12,17), sulprostone had no effect on GSIS in WT or NGOB islets, but reduced GSIS in HGOB islets (Fig. 4B). Interestingly, however, sulprostone reduced the Ca^2+^ duty cycle in all three groups, most strikingly in NGOB and HGOB (Fig. 4C), and significantly reduced cAMP levels in NGOB and HGOB β-cells (Fig. 4D). These results demonstrate that the EP3 receptor is functionally active in all three groups with effects proportional to its relative mRNA expression. Furthermore, the effect of sulprostone in the NGOB islets is the first data to our knowledge demonstrating that a reduction in Ca^2+^ duty cycle and cAMP is not predictive of insulin secretion. Finally, as our Ex4 experiments confirm there is not an overall derangement in β-cell GPCR signaling in NGOB islets, the de-coupling of EP3 signaling from regulation of GSIS may be of biological significance.

**Figure 4.**
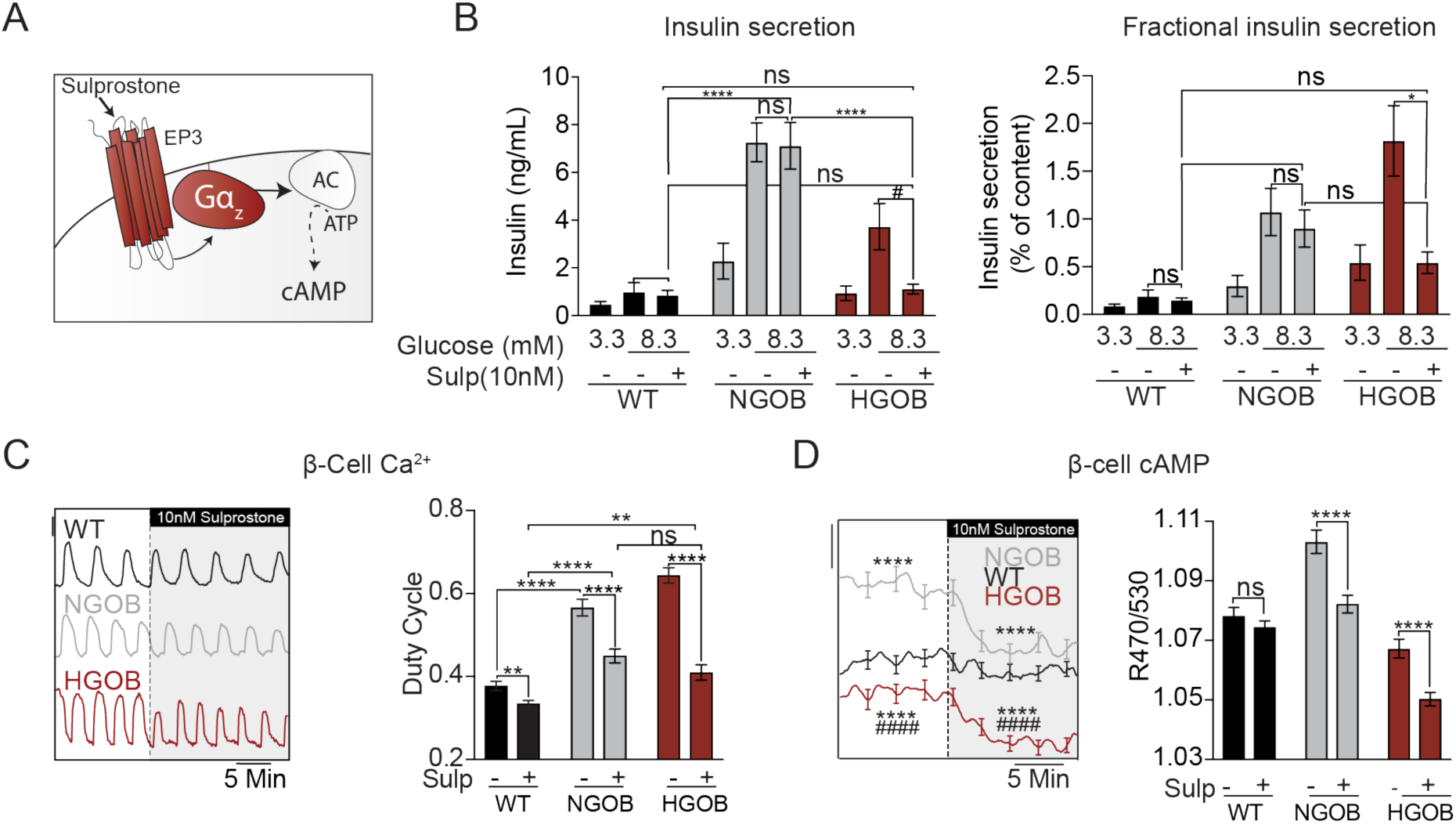
The EP3 agonist sulprostone reduces Ca^2+^ duty cycle and cAMP levels, but inhibits insulin secretion only in failing β-cells. (A) Diagram of the EP3 receptor agonist Sulprostone and its effects on cAMP production (B) Glucose stimulated insulin secretion from WT, NGOB and HGOB islets treated with 3.3 mM glucose or 8.3 mM glucose ± 10 nM sulprostone. Data is represented as both total insulin secreted (left) and fractional insulin secretion as a percent of total insulin content. N=7-10 mice. (C) Representative Ca^2+^ recordings and duty cycle quantification of WT, NGOB and HGOB islets treated with 9 mM glucose followed by 9 mM glucose +10 nM sulprostone. (D) Representative average cAMP recordings and quantification of WT, NGOB and HGOB islets treated with 9 mM glucose followed by 9 mM glucose +10 nM sulprostone. In ‘C’ and ‘D’ N=55-82 islets from 3 mice (NGOB and HGOB) and 84 islets from 6 mice (WT). Scale bar: FuraRed (Ca2+)= 0.01 (B) or 0.025 (C) R470/530 (cAMP). Error bars represent the standard error of the mean (SEM). Data was analyzed by one-way ANOVA or multiple student t-test with Tukey test post-hoc.*p<0.05, **p<0.01, ***p<0.001, ****p<0.0001. For panel D, ****p<0.0001 as compared to WT, #### p<0.0001 as compared to NGOB.

## Discussion

β-cell cAMP is essential for normal cellular function, and impaired β-cell cAMP levels in diabetes pathology have been attributed to alterations in receptor regulated Gα_s_ activity (27,31-36). In this study, we demonstrate a robust increase in cytoplasmic β-cell cAMP and insulin secretion in normoglycemic C57BL/6J *Lep*^*Ob/Ob*^ mice (NGOB) as compared to lean controls that is lost in islets from hyperglycemic C57BL/6J *Lep*^*Ob/Ob*^ (HGOB) mice. β-cell cAMP levels are more predictive of the insulin secretion response than the Ca^2+^ duty cycle, which primarily reflects oxidative glucose metabolism (43,44). Islets from lean, NGOB, and HGOB mice are all sensitive to the incretin analog, exendin-4, to potentiate cAMP levels and GSIS, confirming an intact cAMP amplification pathway. This makes our disparate findings with EP3 expression and downstream signaling in these same islets even more interesting.

PGE_2_ is the most abundant natural ligand of EP3, and is produced from arachidonic acid cleaved from plasma membrane phospholipids by phospholipase A_2_ (PLA_2_) isozymes. The synthesis of arachidonic acid to PGE_2_ is rate-limited by the activity of prostaglandin endoperoxidase synthase (PTGS) 1 and 2 (a.k.a. COX 1 and 2), which catalyze the intermediate conversion of PGH_2_ to PGE_2_. Elevated islet expression of specific PGE_2_ synthetic enzymes and/or islet PGE_2_ excretion has been observed in islets from T2D animals and humans (2,5,6,8,9,20,37). Yet, in this study, PGE_2_ production was not enhanced in HGOB islets above the levels observed in either lean or NGOB. Supporting this, selective EP3 antagonism had no effect on cAMP production or GSIS in HGOB islets. These findings are in contrast to our own previously-published results with islets from *Lep*^*Ob/Ob*^ mice in the BTBR strain background – a very robust model of T2D caused by β-cell dysfunction and loss of functional β-cell mass that occurs with 100% penetrance by 10 weeks of age (2,10). In the B6-OB line, it is possible severe and uncorrected hyperglycemia would, over time, lead to enhanced PGE_2_ synthetic gene expression and up-regulated agonist-dependent EP3 signaling. Yet, the results presented here suggest up-regulation of EP3 itself is the largest contributor to β-cell failure.

The mouse *Ptger3* gene encodes for three splice variants, EP3α, EP3β and EP3γ, all with varying degrees of constitute activity. EP3γ has approximately 80% constitutive activity for coupling to G_i/o_ subfamily proteins (29,45), and it is this splice variant whose expression is sequentially altered in WT, NGOB, and HGOB islets. Not surprisingly, we find EP3γ expression correlates with impaired cAMP levels in HGOB islets. The fact sulprostone significantly reduces NGOB β-cell cAMP and Ca^2+^ levels without significantly affecting GSIS was unexpected, and to our knowledge, is the first report of decoupling second messenger signaling with secretion. Yet, it is the constitutive activity of EP3γ that makes its expression in NGOB islets perplexing. Based on the traditional role of G_i/o_ proteins to block adenylate cyclase activity and reduce cAMP production, EP3γ activity may reduce NGOB β-cell cAMP levels from an even higher theoretical baseline. Numerous signaling changes have been demonstrated in β-cell compensation and failure, such as those mediated by β-cell stress pathways (6,9,38,39), many of which converge on cAMP (14,31,40-42). In this scenario, EP3γ up-regulation may be a compensatory adaptation to dampen β-cell stress and promote β-cell survival. In contrast, molecular signaling mechanisms and their ultimate biological effects may differ in the healthy, compensating, and failing β-cell based on the (patho)physiological milieu in which the islet exists. Therefore, it is possible that the primary effector of constitutively-active EP3γ is not adenylate cyclase, and that shuttling of EP3γ to this alternate effector might even contribute to the elevated cAMP level of NGOB β-cell s. Without an allosteric EP3 antagonist or a β-cell -specific EP3γ B6-OB knockout mouse, these models cannot be differentiated, but will be the subject of future work.

In sum, previous work has implicated the β-cell EP3 receptor as a potential target for T2D therapeutics, but our results with the B6-OB mouse model suggests the contribution of EP3 to β-cell compensation and function is much more nuanced. Future studies should aim to understand the effects of ligand-dependent and -independent EP3 receptor activity in T2D pathology.

## Methods

### Animal Care

C57BL/6J *Lep*^*WT/OB*^ mice were purchased from The Jackson Laboratory and experimental mice bred in-house at the UW-Madison Research Animal Resource Center Breeding Core. All protocols were approved by the Institutional Animal Care and Use Committees of the University of Wisconsin-Madison and the William S. Middleton Memorial Veterans Hospital, which are both accredited by the Association for Assessment and Accreditation of Laboratory Animal Care. All animals were treated in accordance with the standards set forth by the National Institutes of Health Office of Animal Care and Use. Mice were housed in temperature- and humidity-controlled environments with a 12:12-h light/dark cycle and fed pelleted mouse chow (Laboratory Animal Diet 2920; Envigo, Indianapolis, IN) and acidified water (Innovive, San Diego, CA) ad libitum. Only male mice were used in these experiments. Blood glucose was measured by tail prick using an AlphaTRAK glucometer (Zoetis) and rat/mouse test strips. A random-fed blood glucose cut-off of 300 mg/dl was used to define hyperglycemia. Islets were isolated from experimental mice at 10-12 weeks mice utilizing a collagenase digestion procedure as described (47).

### *Ex Vivo* Islet GSIS Assays

GSIS assays were performed 1 day after islet isolation as previously described (48). Briefly, 100 islets were washed and pre-incubated for 45 min in Krebs-Ringer Buffer containing 0.5% BSA and 3.3 mM glucose. Islets were then incubated at a density of 10 islets/ml for an additional 45 min in the indicated treatment. Secretion media was collected and islets were lysed in 1 ml of lysis buffer (20 mM Tris-HCl, pH 7.5; 150 mM NaCl, 1 mM Na_2_EDTA, 1 mM EGTA, 1% Triton X, 2.5 mM sodium pyrophosphate, 1 mM β-glycerophosphate, 1 mM sodium orthovanadate, and 1 μg/ml leupeptin) to determine insulin content. Insulin was assayed by ELISA as previously described (2).

### PGE_2_ Production Assay

On the day of isolation, islets were cultured at a density of 50 islets/ml in normal culture media for 24 hours. Secretion media was collected and cleared by centrifugation for 10 min at 10,000g. Cleared secretion media was stored at −80°C until assayed. PGE_2_ concentrations were determined by monoclonal ELISA (Cayman 515010) following the manufacturers protocol as previously described (2).

### RNA Extraction, cDNA Synthesis, and Transcript Expression Analysis

RNA extraction using TRIzol (ThermoFisher), cDNA synthesis, and real-time quantitative reverse transcriptase PCR analysis (qRT-PCR) was performed using SYBR Green® reagent (Sigma Aldrich) as previously described (46). All gene expression was normalized to that of β-actin. Primer sequences are available upon request.

### Adenovirus Amplification, Purification and Transduction

A plasmid vector encoding the EPAC-based high-affinity cAMP biosensor (Epac-S^H187^) was a kind gift of Dr. Kees Jalink from the Netherlands Cancer Institute (49). This sensor was cloned into an adenoviral vector under the control of the rat insulin promoter in order to generate a β-cell-specific expression construct, as previously described and validated (50). Briefly, H187 was inserted downstream of the rat insulin 1 promoter and rabbit β-globin intron (RIP-BGI). ClonaseII was used to prepare the full-length adenoviral construct in pAd-PL/DEST (Invitrogen), yielding pAd-RIP1-Epac-S^H187^-pA. Adenoviruses were amplified in HEK293 cells and purified by CsCl gradient ultracentrifugation as previously described (27). Immediately after isolation, islets were treated with the high titer virus (0.5 µl/ml) for 2-3 hours and then returned to islet growth medium as previously described (27). All assays utilizing adenovirus were performed 3 days after transduction.

### 2-Photon Imaging of Epac-S^H187^ Expression

Islets expressing pAd-RIP1-Epac-S^H187^-pA were imaged in No. 1.5 glass-bottom dishes on a multiphoton laser scanning system based around a Nikon TE-300 inverted microscope equipped with a 40x/1.15N.A. water immersion objective in a standard imaging solution (135 mM NaCl, 4.8 mM KCl, 5 mM CaCl_2_, 1.2 mM MgCl_2_, 10 mM HEPES, 10 mM glucose; pH 7.35). Temperature was maintained at 37 °C with a Tokai Hit incubator. The Venus fluorescent protein was excited with a Chameleon Ultra laser (Coherent) at 920 nm. Fluorescence emission was captured by a Hamamatsu photomultiplier tube after passing through a 535/70 emission filter (Chroma). Images were collected at 512 × 512 resolution with a 2-µm z-step size at an optical zoom of 1.25 and dwell time of 2 µs.

### Simultaneous Live Cell Islet Ca^2+^ and cAMP Imaging

Wild-type (WT), normoglycemic *Lep*^*Ob*^ (NGOB), or hyperglycemic *Lep*^*Ob*^ (HGOB) islets were imaged simultaneously for islet Ca^2+^ and β-cell cAMP as previously described (27). In each experiment, two groups were imaged, with one group pre-labeled with 0.5 μg/ml DiR (Molecular Probes D12731) for 10 minutes. For measurements of cytosolic Ca^2+^, islets were pre-incubated in 2.5 μM FuraRed (Molecular Probes F3020) in islet media containing 9 mM glucose for 45 min at 37°C. Islets were imaged in a glass-bottomed imaging chamber (Warner Instruments) mounted on a Nikon Ti inverted microscope equipped with a 20x/0.75NA SuperFluor objective (Nikon Instruments). The chamber was perfused with a standard imaging solution (described above). The flow rate was set to 0.25 ml/min by a feedback controller and inline flow meter (Fluigent MCFS EZ). Temperature was maintained at 33°C using solution and chamber heaters (Warner Instruments). Excitation was provided by a SOLA SEII 365 (Lumencor) set to 5-8% output. Single DiR images utilized a Chroma Cy7 cube (710/75x, T760lpxr, 810/90m). Excitation (x) or emission (m) filters (ET type; Chroma Technology) were used in combination with an FF444/521/608-Di01 dichroic beamsplitter (Semrock) as follows: FuraRed, 430/20x and 500/20x, 630/70m (R430/500), for cAMP biosensor FRET imaging, CFP excitation was provided by an ET430/24x filter and emission filters for CFP and Venus emission (ET470/24m and ET535/30m, Chroma) reported as an emission ratio (R470/535). Fluorescence emission was collected with a Hamamatsu ORCA-Flash4.0 V2 Digital CMOS camera every 6 seconds. A single region of interest was used to quantify the average response of each islet using Nikon Elements. Duty cycle analysis was done utilizing custom MATLAB software (Mathworks) as previously described (51).

### Statistical Analyses

Data were analyzed using GraphPad Prism v. 8 (GraphPad Software Inc., San Diego, CA). A (1) T-test or (2) one-way ANOVA, two-way ANOVA analysis followed by Tukey post-hoc correction for multiple comparisons was used as appropriate. P< 0.05 was considered statistically significant.

## Acknowledgements

The authors wish to thank Kathryn Carbajal, Trillian Gregg, Helena VanDeusen, Steve Trier, and Melissa Skala for their scientific and technical guidance during this work. This work was supported by Merit Review Award I01 BX003700 (to M.E.K.) from the United States (U.S.) Department of Veterans Affairs Biomedical Laboratory Research and Development (BLR&D) Service. This work was also supported in part by National Institutes of Health (NIH) grant R01 DK102598 (to M.E.K.), R01 DK113103 (to M.J.M.), and R01 AG062328 (to M.J.M.); American Diabetes Association Grant 1-14-BS-115 (to M.E.K.); and start-up and pilot project funding from the University of Wisconsin-Madison Department of Medicine, Graduate School, and Office of the Provost (to M.E.K. and M.J.M). The study sponsors had no role in the study design; collection, analysis or interpretation of data; the writing of the report; or the decision to submit the paper for publication. The content is solely the responsibility of the authors and does not necessarily represent the official views of the National Institutes of Health, the U.S. Department of Veterans Affairs, or the United States Government.

## Author Contributions Statement

MDS, MJM, and MEK conceived of and designed the work. MDS, JMH, GMK, and SMS acquired and analyzed the data. MDS, MJM, and MEK interpreted the data. MDS drafted the manuscript and the figures, and MDS and MEK revised them. All authors reviewed the manuscript.

